# Neural crest-related NXPH1/α-NRXN signaling opposes neuroblastoma malignancy by inhibiting metastasis

**DOI:** 10.1101/2021.11.26.470092

**Authors:** Lucía Fanlo-Escudero, Soledad Gómez-González, Irene Sangrador, Emmanuel L. Gautier, Susana Usieto, Elena Rebollo, Mònica Vila-Ubach, Ángel M. Carcaboso, Toni Celià-Terrassa, Cinzia Lavarino, Elisa Martí, Gwenvael Le Dréau

## Abstract

Neuroblastoma is a pediatric cancer that can present as low- or high-risk tumors (LR-NBs and HR-NBs), the latter group showing poor prognosis due to metastasis and strong resistance to current therapy. NBs are known to originate from alterations to cells in the sympatho-adrenal lineage derived from the neural crest, but whether LR-NBs and HR-NBs differ in the way they exploit the transcriptional program underlying their developmental origin remains unclear. Here, we compared the transcriptional landscapes of primary samples of LR-NBs, HR-NBs and human fetal adrenal gland, and thereby identified the transcriptional signature associated to NB formation that further distinguishes LR-NBs from HR-NBs. The majority of the genes comprising this signature belong to the core sympatho-adrenal developmental program, are associated with favorable patient prognosis and with diminished disease progression. The top candidate gene of this list, *Neurexophilin-1 (NXPH1)*, encodes a ligand of the transmembrane receptors *α-Neurexins* (*α-NRXNs*). Our functional *in vivo* and *in vitro* assays reveal that NXPH1/α-NRXN signaling has a dual impact on NB behavior: whereas NXPH1 and α-NRXN1 promote NB tumor growth by stimulating cell proliferation, they conversely inhibit the ability of NB cells to form metastases. Our findings uncover a module of the neural crest-derived sympatho-adrenal developmental program that opposes neuroblastoma malignancy by impeding metastasis, and pinpoint NXPH1/α-NRXN signaling as a promising target to treat HR-NBs.

## Introduction

Neuroblastoma (NB) represents the most common cancer in infants and the most frequent solid extracranial tumor in childhood, accounting for 10-15% of cancer-related child deaths (1–3). NBs are nearly all sporadic (98%) and are very heterogeneous (4), half of the tumors being detected in para-spinal ganglia in the abdomen, chest and neck, while the other half are found in the adrenal gland medulla (5). Some 60% of NBs are classified as low-to-intermediate risk (LR-NBs) and correspond mostly to loco-regional tumors that present a good prognosis and can even regress spontaneously (6, 7). By contrast, the remaining 40% are classified as high-risk tumors (HR-NBs) and are often already metastatic at the time of diagnosis, displaying a strong resistance to therapy and a high probability of relapse that is reflected in poor patient outcome (8, 9). The most predictive genetic events of a poor prognosis for NB patients appear to be amplification of the proto-oncogene *MYCN*, and mutations in the *ALK, ATRX* and *PHOX2B* genes (6,8,10). However, these genetic alterations are only found in up to 16% of all NB cases and thus, they are insufficient to explain the differences in the aetiology and malignancy of HR-NBs relative to LR-NBs (4–6). Accordingly, the challenge remains to identify the genetic and molecular mechanisms that discriminate LR-NBs from HR-NBs.

Lately, much attention has been oriented towards the identification of the NB cell-of-origin, with the goal to determine whether LR-NBs and HR-NBs differ in the way they exploit the transcriptional program of this cell-of-origin. There is a consensus on the fact that NBs originate from alterations to cells in the sympatho-adrenal (SA) lineage (11–16). Representing one of the multiple cell fate branches of neural crest cells, the SA lineage is composed of four main cell identities: Schwann cell precursors, bridge cells, chromaffin cells and sympathoblasts (12–17). The specific identity of the NB cell-of-origin remains however debated (11–16). Thus, more work is needed to elucidate whether LR-NBs and HR-NBs differ in the way they exploit the transcriptional program underlying their developmental SA origin, and even more importantly, if this SA program could be exploited to identify novel factors capable of impeding the metastatic dissemination and malignancy of HR-NBs.

Here, we addressed these questions by first comparing the genome-wide transcriptomic profiles of primary samples from LR-NB and HR-NB patients and human fetal adrenal gland, and thereby identified a transcriptional signature associated to NB formation that further distinguishes LR-NBs from HR-NBs. We then established that LR-NBs and HR-NBs can be discriminated by a signature corresponding to the core SA lineage, rather than by signatures specific of the distinct SA cell identities. Remarkably, this core SA signature was composed in vast majority of genes associated with a favorable patient prognosis and with diminished disease progression. From these findings we selected *Neurexophilin-1*, which encodes the secreted glycoprotein NXPH1, as a candidate for functional studies and demonstrated that its activity, mediated by its receptors α-NRXN1/2, indeed represses NB malignancy by inhibiting the ability of NB cells to form metastases.

## Results

### NB formation is correlated with a neural signature accentuated in LR-NBs

We set out to define the transcriptional signatures associated to the aetiology of LR- and HR-NBs and to the malignant behavior of HR-NBs. As such, we analyzed the transcriptome of primary tumor samples from 18 NB patients. These samples were classified as LR (n=8) or HR (n=10) based on their tumor stage (International Neuroblastoma Staging System INSS 1-3, 4s and 4) (31), their *MYCN* amplification status and their age at diagnosis (Fig. 1A, Supplementary Fig.S1A and Supplementary Table S1) (20). Samples of the human fetal adrenal gland (fAG) were used as a normal reference tissue (Fig. 1A, Supplementary Fig.S1A and Supplementary Table S1).

**Figure 1:**
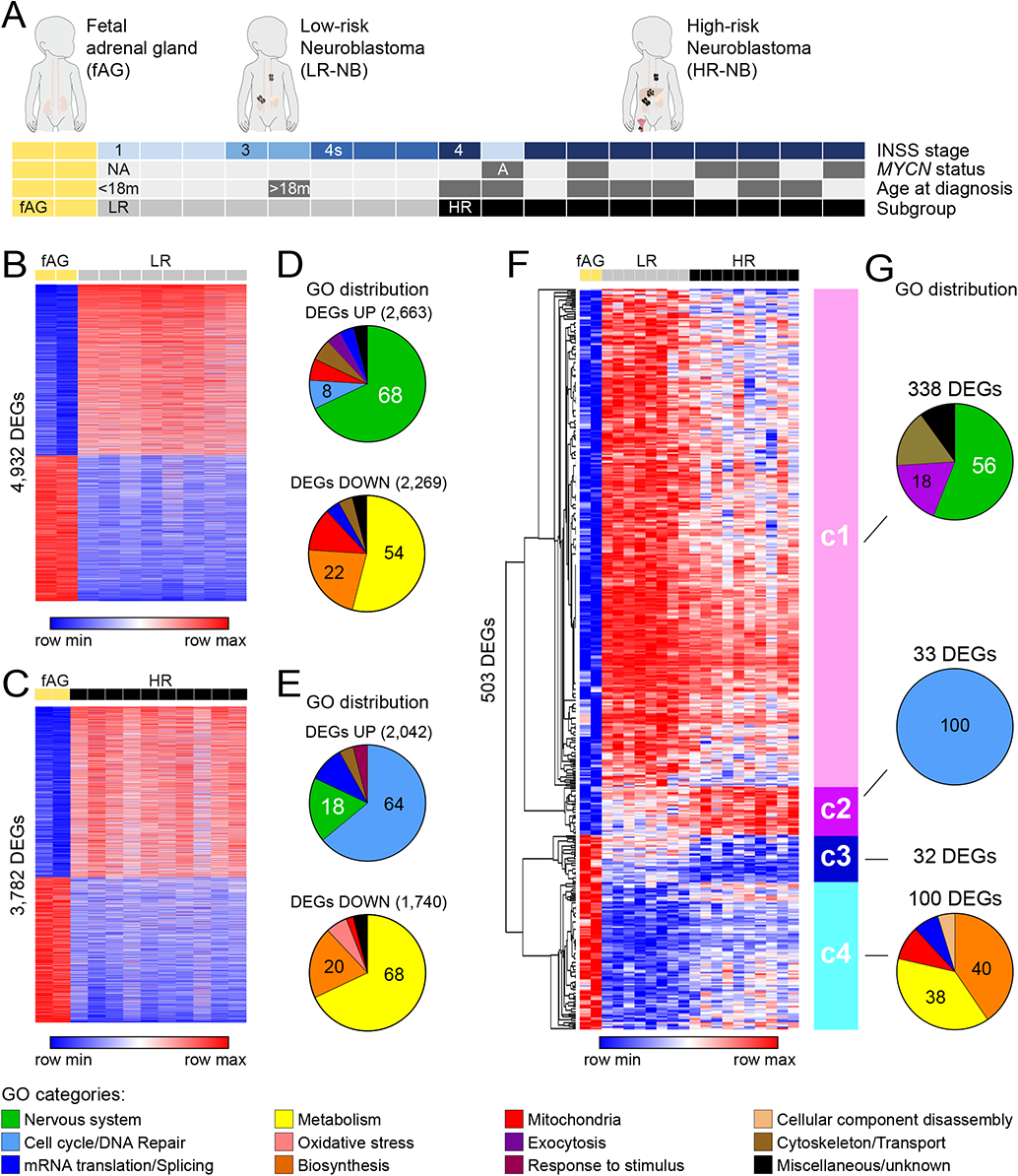
NB formation is correlated with a neural signature accentuated in LR-NBs. (**A**) The primary samples of human neuroblastoma (NB) patients were classified as low-risk (LR, n=8) or high-risk (HR, n=10) based on their *MYCN* amplification status (A: amplified or NA: non-amplified), the INSS tumor stage (1-3, loco-regional; 4s, metastatic but regressing; 4, metastatic and therapy-resistant) and the age at diagnosis (< or >18 months). Samples of human fetal adrenal gland were used as a healthy reference tissue (fAG, n=2). (**B, C**) Heatmap representations of the differentially expressed genes (DEGs) obtained by comparing (**B**) LR *vs* fAG and (**C**) HR *vs* fAG, using a LIMMA R package (adjusted P<0.05). (**D, E**) Distribution of the 50 most enriched gene ontology (GO) biological processes grouped by ancestor annotation for the upregulated (upper panel) and downregulated (lower panel) lists of DEGs from the (**D**) LR *vs* fAG and (**E**) HR *vs* fAG comparisons. (**F**) Heatmap of the 503 DEGs identified by comparing HR*vs*LR, using the list of 3,096 DEGs common to LR*vs*fAG and HR*vs*fAG. An unbiased hierarchical clustering analysis subdivided these DEGs into 4 clusters (c1-c4). (**G**) Distribution of the most enriched (up to 50) GO biological processes grouped by ancestor annotation for the clusters c1,c2 and c4 (no GO enrichment for c3).

To identify the transcriptional signatures associated to NB aetiology, we first independently compared the LR- and HR-NB groups to the fAG samples and thereby identified 4,932 and 3,782 differentially expressed genes (DEGs), respectively (Fig. 1B, C and Supplementary Table S1). A gene ontology (GO) enrichment analysis revealed that the downregulated DEGs retrieved from the LR *vs* fAG and HR *vs* fAG comparisons were associated mainly with GO terms related to metabolism and biosynthesis (Fig. 1D, E and Supplementary Table S2), in agreement with previous findings (11). By contrast, the DEGs found upregulated in these two comparisons were both characterized by a marked enrichment in GO terms related to the nervous system, or to the cell cycle and DNA repair (Fig. 1D, E and Supplementary Table S2). The GO terms related to the nervous system represented 68% of the top 50 GOs for the LR *vs* fAG comparison but only 18% for the HR *vs* fAG comparison (Fig. 1D, E and Supplementary Table S2), indicating that the formation of both LR- and HR-NBs is associated to a neural signature, and that this signature is more pronounced in LR-NBs.

We next searched for a transcriptional signature relevant to NB formation that furthermore distinguishes LR-NBs from HR-NBs. To this aim, we crossed the two independent comparisons LR *vs* fAG and HR *vs* fAG and thereby retrieved 3,096 common DEGs, which included the well-established markers of NB malignancy: *ALK*, *ATRX*, *MYCN*, *PHOX2A* and *PHOX2B* (Supplementary Fig. S1B-D and Supplementary Table S3). We then searched among these 3,096 common DEGs for the genes that were further differentially expressed between the LR-NB and HR-NB groups. This comparison identified a list of 503 LR *vs* HR DEGs that an unbiased hierarchical clustering analysis subdivided into 4 clusters of genes presenting distinct expression profiles (Fig. 1F and Supplementary Table S3). The clusters c1 and c2 consisted of genes more strongly expressed in both LR-NBs and HR-NBs than in fAG samples. The best represented cluster (c1) contained 338 genes that were more strongly expressed in LR-NBs than in HR-NBs and were associated with a marked enrichment in GO terms related to the nervous system (Fig. 1F, G and Supplementary Tables S3 and S4). The expression levels of the top 30 genes found in cluster c1 (such as *SOX6*, *NXPH1*, *PRPH*) were all correlated with a favorable patient prognosis, as shown by Kaplan-Meyer analyses done with an independent cohort of 498 NB patients (Supplementary Fig. S1E and Supplementary Table S3). Conversely, the 33 genes found in cluster c2 (such as *CDCA7*, *C4orf46*, *THOC4*) were more strongly expressed in HR-NBs than in LR-NBs and presented the features of oncogenes, as shown by their GO term enrichment profile and Kaplan-Meyer analyses (Fig. 1F, G, Supplementary Fig. S1E and Supplementary Tables S3 and S4). On the other hand, the clusters c3 and c4 consisted of genes expressed more weakly in both LR-NBs and HR-NBs than in fAG samples. The 32 genes found in cluster c3 presented the features of tumor suppressors, while the cluster c4 contained 100 genes that were more expressed in HR-NBs than in LR-NBs and were mainly correlated with a bad prognosis (Fig. 1F, G, Supplementary Fig. S1E and Supplementary Tables S3 and S4). Our analysis thus identified a complex transcriptional signature relevant to NB formation that furthermore distinguishes LR-NBs from HR-NBs. Remarkably, 67% of the 503 genes found in this signature, which formed the cluster c1, are both associated with a neural identity and a better patient outcome.

### The core sympatho-adrenal signature is enriched in LR-NBs and associates with better patient prognosis

We next assessed whether the transcriptional signature distinguishing LR-NBs and HR-NBs was related to their sympatho-adrenal (SA) origin. The human SA lineage can be subdivided into 4 main cell identities: Schwann cell precurors (SCPs), bridge cells, chromaffin cells and sympathoblasts (Fig. 2A) (12–16). We searched for a possible enrichment of these SA cell identities within our list of LR *vs* HR DEGs using transcriptional signatures recently identified by single-cell RNA-seq during human adrenal gland development (16). We thereby observed that the 4 SA cell signatures all showed a strong enrichment for genes expressed at higher levels in LR-NBs than in HR-NBs, retrieving as many as 342 of our 503 DEGs (Fig. 2B and Supplementary Table S5). By contrast, the signatures of the non-adrenal medulla cell types identified in the developing human adrenal gland did not show any enrichment, except the adrenal cortex signature which was enriched in genes expressed at higher levels in HR-NBs (Supplementary Fig. S2A and Supplementary Table S5). We noticed that the full SCP, bridge cell, chromaffin cell and sympathoblast signatures were largely overlapping (Fig. 2C and Supplementary Table S5). We retrieved a strong enrichment for genes expressed at higher levels in LR-NBs when we used a core SA signature consisting of the 4,518 genes shared by at least 3 of the 4 SA cell signatures (Fig. 2D and Supplementary Table S5). By contrast, this enrichment was lost when using the signatures specifically consisting of the genes unique to SCPs, bridge cells or sympathoblasts, and the “unique” chromaffin cell signature showed a converse and weak enrichment towards genes expressed at higher levels in HR-NBs (Supplementary Fig. S2B, C). Using the core SA signature we retrieved more than half of the 503 LR *vs* HR DEGs (Fig. 2D and Supplementary Table S6). Even more strikingly, 92% of this list (242 genes out of 262 genes) corresponded to genes belonging to the cluster c1, thereby representing 72% of this whole cluster c1 (Fig. 2E and Supplementary Table S6). The transcriptional landscapes of LR-NB and HR-NBs can thus be discriminated based on a signature largely corresponding to the core SA lineage, and this developmental program is unexpectedly associated with a favorable prognosis.

**Figure 2:**
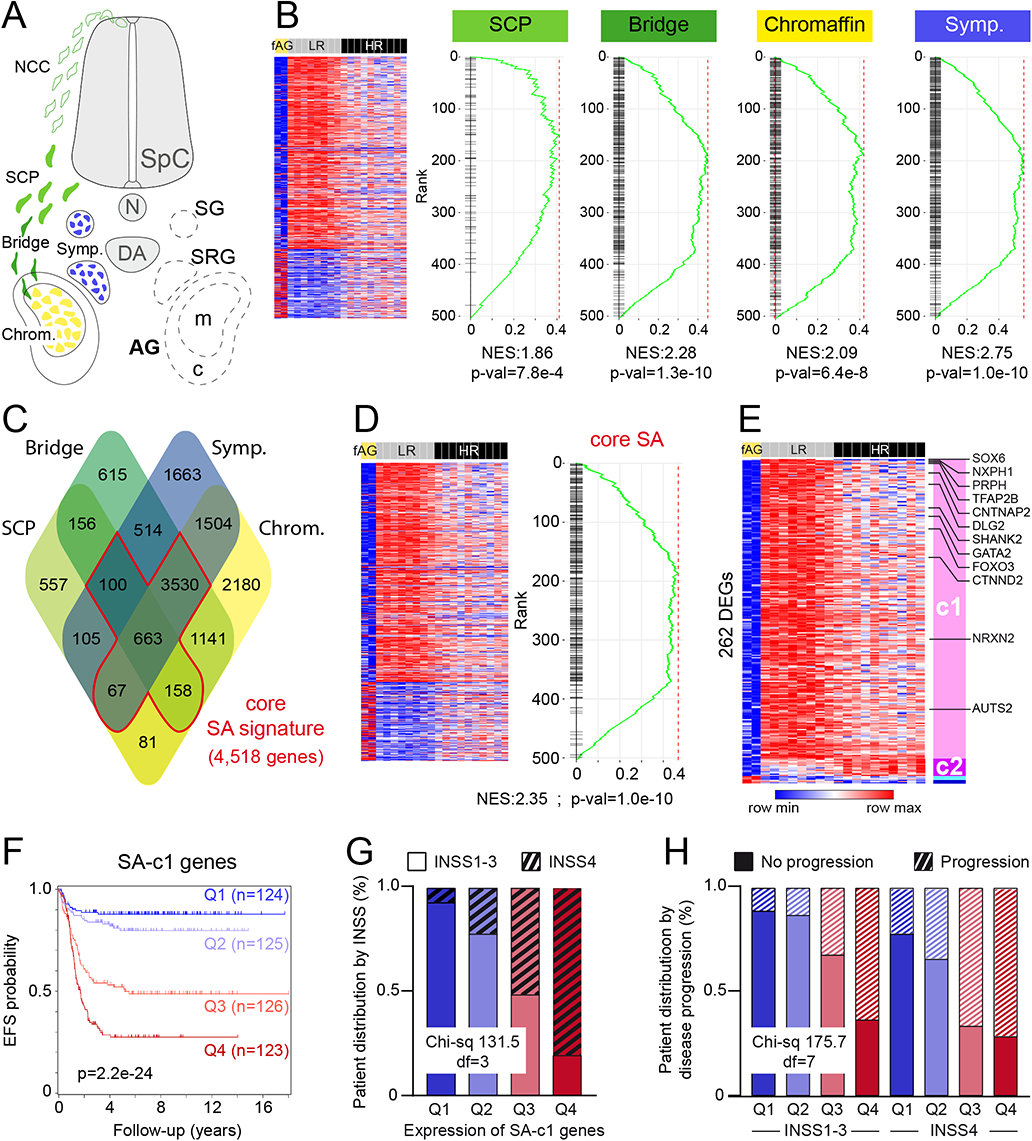
The core sympatho-adrenal signature is enriched in LR-NBs and associates with better patient prognosis. (**A**) Schematic representation of the 4 cell identities forming the neural crest-derived sympatho-adrenal lineage, including Schwann cell precursors (SCPs, light green), bridge cells (dark green), chromaffin cells (yellow) and sympathoblasts (blue). (**B**) Gene set enrichment analysis (GSEA) of the signatures of human SCPs, bridge cells, chromaffin cells and sympathoblasts within the list of HR *vs* LR DEGs. (**C**) Venn diagram showing the overlapping of the human SCP, bridge cell, chromaffin cell and sympathoblast signatures, highlighting its core sympatho-adrenal (SA) signature. (**D**) GSEA of the core SA signature within the list of HR *vs* LR DEGs. (**E**) Heatmap of the 262 HR *vs* LR DEGs retrieved in the core SA signature, organised by clusters and fold changes. (**F**) Follow-up of the event-free survival (EFS) probability of a cohort of 498 NB patients (SEQC database, R2 online platform) based on the expression of the 242 SA-c1 genes, the cohort being subdivided into 4 quartiles (Q1-Q4, from higher to lower numbers of SA-c1 genes showing expression levels above average). (**G**) Distribution of the SA-c1 quartiles based on INSS tumor stage (loco-regional stages 1-3, lower filled bars; metastatic stage 4, upper cross-hatched bars). (**H**) Distribution of the 8 SEQC sub-groups identified in (G, INSS1-3 or INSS4 for each of the SA-c1 quartiles Q1 to Q4) according to the absence of presence of disease progression. Significance was automatically assessed by the R2 server using log-rank test (**F**) or with the Chi-square and Fisher’s tests (**G, H**). AG, Adrenal gland; c, adrenal cortex; DA, dorsal aorta; m, adrenal medulla; NCCs, neural crest cells; N, notochord; SpC, spinal cord; SG, sympathetic ganglia; SRG, supra-renal ganglia; NES, normalized enrichment score.

Among these 242 genes common to the core SA signature and to the LR *vs* HR DEGs of the cluster c1 (thereafter named “SA-c1” genes) were found for instance the transcription factors (TFs) SOX6 and TFAP2 involved in the early stages of the NCC lineage (12, 32), PRPH and the TFs GATA2 and FOXO3, which are associated with chromaffin cell differentiation (12), and various autism spectrum disorder genes (*CNTNAP2*, *CTNND2, DLG2*, *NRXN2* and *SHANK2*) recently associated with NB aetiology (33) (Fig. 2E and Supplementary Table S6). To test the predictive potential of the SA-c1 gene module on NB prognosis, we subdivided the SEQC NB cohort into quartiles based on the combined expression of the 242 SA-c1 genes, whereby the quartiles Q1 and Q4 consisted of the patient samples presenting the highest and lowest numbers of SA-c1 genes expressed above average levels, respectively (see Materials and Methods). These quartiles showed a progressive decrease in event-free survival probability (8-years EFS probability of 0.88, 0.80, 0.49 and 0.27 for Q1, Q2, Q3 and Q4, respectively, Fig. 2F). They also consisted of increasing proportions of INSS4 tumors, which are metastatic and aggressive (31), at the expense of INSS1-3 tumors usually associated with a good prognosis (Fig. 2G). Importantly, the positive correlation between SA-c1 gene expression and better prognosis was also observed when focusing on either INSS1-3 or INSS4 tumors (Supplementary Fig. S3A, B). Remarkably, the expression levels of the SA-c1 module correlated to patient prognosis more strongly than the INSS classification. For instance, the group of INSS4 samples from the quartile Q1 presented a higher EFS probability than the groups of INSS1-3 samples from the quartiles Q3 and Q4 (Supplementary Fig. S3A, B). We further sub-divided the SEQC cohort based on the progression of the disease, including within the category “progression” the tumors that did not respond to therapy plus those that were recurrent despite an initial therapeutic response. We thereby observed an inverse correlation between disease progression and the combined expression of the SA-c1 genes (Fig. 2H). The module of SA-c1 genes related to the core sympatho-adrenal program is thus strikingly predictive of NB malignancy, patient prognosis and disease progression. The SA-c1 genes might thus play a crucial role in restraining NB malignancy.

### The expression of *NXPH1* and its receptors *α-NRXN1/2* associates with favorable patient prognosis and identifies NB cells with a neural crest stem cell identity

To functionally test the impact of SA-c1 genes on NB malignancy, we selected *Neurexophilin-1* (*NXPH1*). *NXPH1* was the SA-c1 gene with the second highest fold change enrichment in LR-NBs relative to HR-NBs (fc LR/HR=7.46; Fig. 2E and Supplementary Table S6) and its expression levels strongly correlated with favorable patient prognosis and with diminished disease progression in both the INSS1-3 and INSS4 NB sub-groups (Fig. 3A, B and Supplementary Fig. S3C-E). *NXPH1* is mostly expressed in the nervous system and encodes a secreted glycoprotein that specifically binds to and modulates the activity of the α-Neurexin transmembrane receptors (α-NRXN1, 2 and 3; Fig. 3C), which are known to play key roles in synaptogenesis and neurotransmission (34–37). Interestingly, *NRXN2* also came out as a SA-c1 gene (Fig. 2E and Supplementary Table S6) and both *NRXN1* and *NRXN2* expression levels associated with a favorable prognosis, whereas *NRXN3* levels did not (Fig. 3D and Supplementary Fig. S3F).

**Figure 3:**
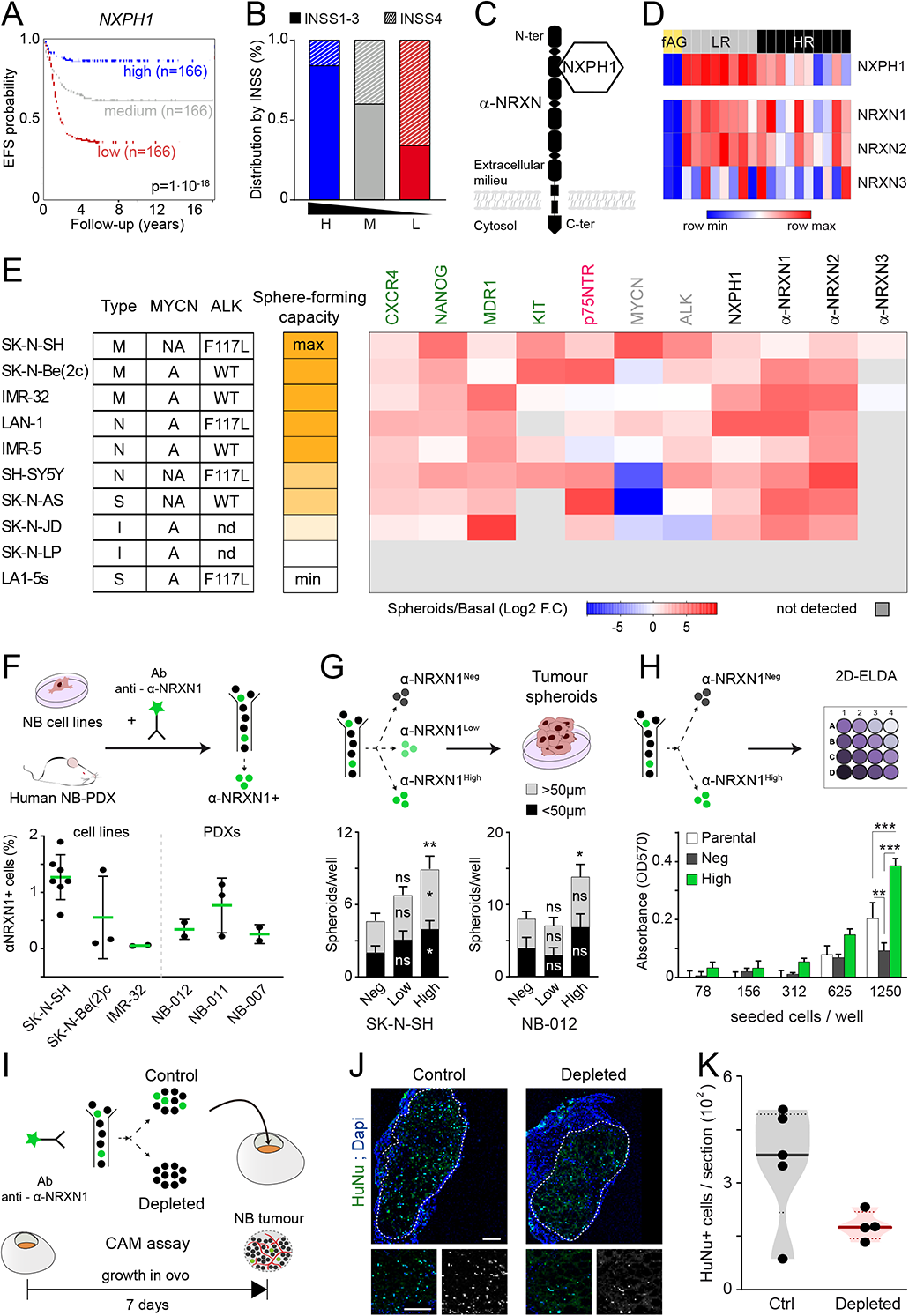
The expression of *NXPH1* and its receptors *α-NRXN1/2* associates with favorable patient prognosis and identifies NB cells with a neural crest stem cell identity. (A) EFS probability and (B) distribution of the SEQC cohort subdivided into 3 groups based on NXPH1 transcript levels (high, medium or low, A, B) and INSS tumor stage (stages 1-3, filled bars; stage 4, dashed bars, B). (C) Schematic representation of NXPH1 and its α-NRXN receptors. (D) Heatmap of NXPH1 and NRXN1-3 expression in our cohort of fAG, LR-NB and HR-NB samples. Note that only NXPH1 and NRXN2 belong to the SA-c1 gene set. (E) Heatmap of the transcript levels of stem cell markers (green), a neural crest cell marker (pink), pathogenic NB markers (gray), NXPH1 and α-NRXN1-3 (black), quantified by RT-qPCR in 10 human NB cell lines with different morphologies (M, mixed; N, neuronal; I, intermediate; S, stromal), MYCN status (A: amplified; NA: non-amplified), ALK status (activating mutation F117L; WT: wild-type; nd: not determined) and sphere-forming capacity, after 5 weeks in sphere-forming culture conditions relative to their levels in basal culture conditions. The values were calculated from 2 independent experiments. (F) Mean percentage (± s.d.) of α-NRXN1+ cells in 3 NB cell lines and 3 PDX samples purified by FACS using an anti-α-NRXN1 antibody conjugated to Alexa-488. (G) Mean number (± s.e.m.) of spheroids with a diameter < or > 50 µm generated by FACS-purified α-NRXN1-, α-NRXN1+^Low^ and α- NRXN1+^High^ cells from the SK-N-SH cell line or the PDX NB-012. The values represent the mean of 4 biological (SK-N-SH) or technical (NB PDX) replicates. (H) Extreme limiting dilution assay performed in vitro (2D-ELDA) with non-purified parental SK-N-SH cells (white bars), or FACS-purified SK-N-SH α-NRXN1-(dark grey bars) and α-NRXN1+^High^ (green bars) cells. The mean absorbance at 570 nm (± s.e.m.) was used as an indicator of cell density and quantified 2 weeks after seeding. The values represent the mean of 3 biological replicates. (I) Representation of the chick chorio-allantoid membrane (CAM) assay performed to compare the growth potential of FACS-purified SK-N-SH cells deprived from their α-NRXN1+ cell sub-population (depleted) with that of non-deprived cells (control, Ctrl). (J) Representative images of the tumors formed by HuNu+ (green) NB cells and (K) mean numbers of HuNu+ cells (± s.e.m.) quantified per tumor section 7 days after seeding α-NRXN1+-depleted (n=4) or control (n=5) SK-N-SH cells onto the CAM of E10 chicken embyos. Each dot represents the mean value of an individual tumor calculated from 6-12 images. Significance was assessed using (A) a log-rank test, (B) the Chi-square and Fisher’s tests, (G) a non-parametric Kruskal-Wallis test and a Dunn’s post hoc test or (H) a two-way ANOVA and a post hoc Tukey’s test: *p<0.05; **p<0.01; ***p<0.001; ns, p>0.05. Scale bar: 100μm.

We first characterized the expression of *NXPH1* and its *α-NRXN1/2/3* receptors in a panel of 10 human NB cell lines with diverse genetic profiles and morphological properties (Fig. 3E) (19, 38). In basal culture conditions *NXPH1* and *α-NRXN1/2* were expressed weakly, while the expression of *α-NRXN3* transcripts was below the threshold of detection. Growing NB cell lines in restrictive culture conditions (see Materials and Methods) for 5 weeks *in vitro* can lead to the formation of spheroids that are progressively enriched in cells presenting stem cell-like characteristics and a more specific NCC identity (39–41). Following this protocol, most of the NB cell lines (8 out of 10) showed the capacity to grow as spheroids (Fig. 3E). The formation of spheroids correlated to a stem cell-enrichment, as shown by increased levels of the transcripts encoding the stereotypic stem cell markers *CXCR4*, *NANOG* and *KIT*, the NCC stem cell-specific marker *p75NTR* and the *MDR1* gene associated with chemotherapy resistance (Fig. 3E) (42–44). Remarkably, *NXPH1* and *α-NRXN1/2* levels increased in all the NB cell lines harbouring a sphere-forming capacity (Fig. 3E), thereby revealing a strong positive correlation between the expression of *NXPH1* and *α-NRXN1/2* and the acquisition of a NCC stem cell identity.

Taking advantage of a commercial fluorescence-conjugated antibody recognizing the extracellular region of human α-NRXN1, we identified a small subpopulation of α-NRXN1^+^ cells (less than 1.5%) within the 3 human NB cell lines that showed the highest sphere-forming capacity and in cells dissociated from 3 different NB patient-derived xenografts (PDX; Fig. 3F). Purifying α-NRXN1^high^ cells from the SK-N-SH cell line or from the PDX NB-012 by FACS and growing them in restrictive culture conditions for a week revealed their increased spheroid-forming capacity compared to both α-NRXN1^low^ and α-NRXN1^-^ cells (Fig. 3G). In addition, α-NRXN1^high^ cells harboured an enhanced proliferative ability when seeded under conditions of extreme dilution, relative to purified α-NRXN1^-^ cells or unsorted SK-N-SH cells (Fig. 3H). To assess the importance of α-NRXN1^+^ cells *in vivo*, we compared the growth potential of SK-N-SH cells deprived of their α-NRXN1^+^ cell subpopulation with that of non-deprived cells using a CAM assay, in which cells were seeded on the richly vascularised chorio-allantoid membrane of 10 days-old chicken embryos (Fig. 3I). After 7 days of incubation *in ovo*, the number of cells quantified per tumor section was decreased by ∼50% for the α-NRXN1^+^-deprived cells relative to their control (Fig. 3J, K), thus revealing that α-NRXN1^+^ cells are required to support NB tumor growth *in vivo*. These findings established a correlation between *NXPH1/α-NRXN* expression and the tumorigenic potential of NB cells, arguing that NXPH1/α-NRXN signaling could control NB growth and/or aggressiveness.

### NXPH1/α-NRXN signaling stimulates NB growth

To study whether NXPH1/α-NRXN signaling regulates NB growth, we generated stable clones of SK-N-SH cells that constitutively express sh-RNAs targeting either *NXPH1* or *α-NRXN1* (Supplementary Fig. S4A-D). Both *NXPH1* and *α-NRXN1* knockdowns severely decreased cell viability over a 3 days-period *in vitro* and caused a complete growth arrest after one week (Supplementary Fig. S4E, F). To circumvent this effect, we generated clones whose sh-RNA and eGFP production was only induced upon doxycycline treatment (Supplementary Fig. S4G). Using eGFP^+^ FACS-purified cells pre-treated with doxycyline for 96 hours, we observed that the growth arrest caused by both *NXPH1* and *NRXN1* knockdown was minimized in the inducible sh-NXPH1 and sh-αNRXN1 clones when grown in basal culture conditions (Supplementary Fig. S4H-O). These sh-NXPH1 and sh-αNRXN1 cells however retained a diminished sphere-forming capacity when grown in restrictive culture conditions (Fig. 4A-C). We next tested the effect of these inducible sh-NXPH1 and sh-αNRXN1 clones on NB growth *in vivo*, by xenografting FACS-purified, doxycyline-treated cells into the flanks of immuno-compromised NOD/SCID mice that received doxycycline *ad libitum* (Fig. 4A). The sh-NXPH1 and sh-αNRXN1 cells produced fewer and smaller tumors than their control and caused a marked delay in the time required for their detection (Fig. 4D, E). Thus, inhibiting NXPH1/α-NRXN1 signaling strongly impairs NB tumor formation *in vivo*.

**Figure 4:**
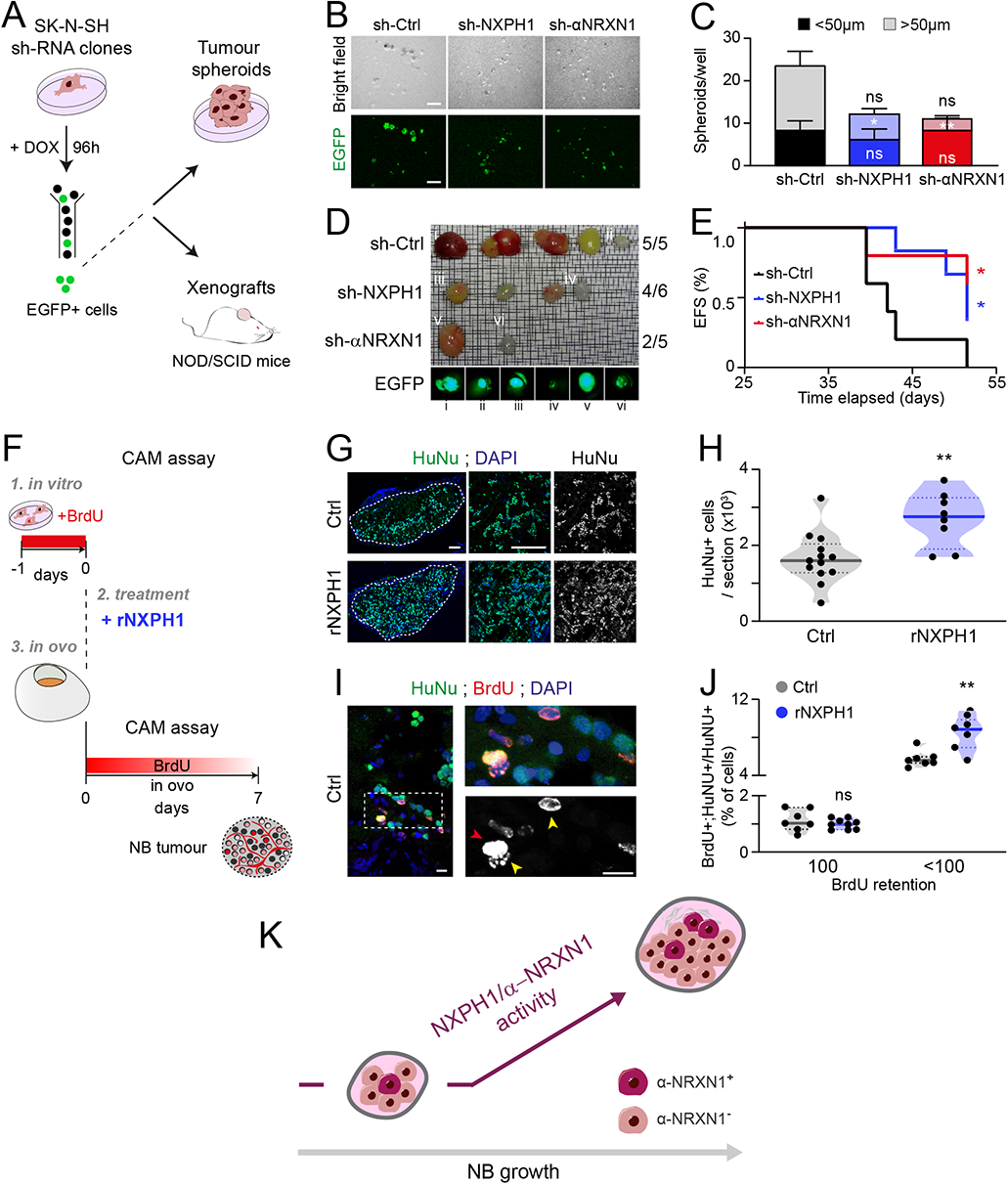
The activity of NXPH1 and α-NRXN1 is required for NB growth. (**A**) The effects of inducible sh-NXPH1 and sh-αNRXN1 clones on NB growth was tested (**B, C**) after xenografting and (**D, E**) *in vitro.* (**B**) Representative images and (**C**) mean numbers of spheroids (± s.e.m.) generated by pre-activated and FACS-purified sh-Ctrl, sh-NXPH1 and sh-αNRXN1 cells grown for 2 weeks under restrictive culture conditions. Spheroids with a diameter < or > 50 µm are indicated. The values represent the result of 8 technical replicates obtained from 2 independent experiments. (**D**) The number and volume of the tumors generated by pre-activated (via doxycyline treatment for 96 hours *in vitro*, +DOX) and FACS-purified sh-Ctrl, sh-NXPH1 and sh-αNRXN1 cells 53 days after xenografting into the flanks of immuno-compromised NOD/SCID mice. (**E**) EFS follow-up of mice xenografted with sh-Ctrl (black, n=5), sh-NXPH1 (blue, n=6) and sh-αNRXN1 (red, n=5) cells. (**F**) Experimental design of the CAM assay used to assess the effects of human recombinant NXPH1 (rNXPH1) on NB cell growth. (**G**) Representative sections of tumors formed by HuNu^+^ (green) NB cells 7 days after seeding onto the CAM of E10 chicken embryos. (**H**) Mean numbers of HuNu^+^ cells (± s.e.m.) quantified per tumor section for the SK-N-SH cells seeded onto Matrigel in presence of 10 µg/ml rNXPH1 (n=8 tumors) or its BSA vehicle (Ctrl, n=13 tumors). Each dot represents the value of an individual tumor calculated from 3-6 images. (**I**) Representative image showing HuNu^+^ cells with complete (100%, red arrowhead) or diluted (<100%, yellow arrowhead) BrdU retention. (**J**) Mean percentage of BrdU^+^; HuNu^+^/total HuNu^+^ cells (± s.e.m.) with 100% or <100% BrdU retention for the SK-N-SH cells seeded onto Matrigel in presence of 10 µg/ml rNXPH1 (n=9 tumors) or its vehicle (Ctrl, n=7 tumors). Each dot represents the value of an individual tumor calculated from 2-3 images. (**K**) Scheme proposing that the increased expression of NXPH1 accompanying NB formation stimulates NB growth by activating α-NRXN1 signaling. Significance was assessed with a two-sided unpaired t-test (**H**), a non-parametric Mann-Whitney test (**J**), a Mantel-Cox log-rank test (**E**) or a Kruskal-Wallis test with a *post hoc* Dunn’s test (**c**): *p<0.05; **p<0.01; ns, p>0.05. Scale bar: 10 μm (**I**), 50 μm (**B, G**).

We next wondered whether stimulating NXPH1/α-NRXN signaling would be sufficient to increase the growth of NB tumors. To test this idea, we opted for a pharmacological gain-of-function approach and performed a CAM assay in which parental SK-N-SH cells were embedded in Matrigel in the presence or absence of recombinant human NXPH1 (rNXPH1, 10 µg/ml; Fig. 4F). After 7 days, we found that rNXPH1 treatment had increased the number of NB cells per tumor section by 62% (Fig. 4G, H and Supplementary Fig. S5A-D). As SK-N-SH cells were incubated with BrdU for 24 hours *in vitro* before seeding on the CAM, we could moreover determine that rNPHX1 addition increased the fraction of NB cells that actively cycled during the incubation *in ovo* (BrdU-ir<100%; Fig. 4I, J). By contrast, rNPHX1 treatment did not affect the proportion of NB cells that became quiescent or cycled very slowly during that period (BrdU-ir=100; Fig. 4I, J), nor apoptosis (Supplementary Fig. S5E, F). Interestingly, the enhanced proliferation rate caused by rNPHX1 was associated with a 2-fold increase in the proportion of NB cells expressing the neural crest stem cell marker p75/NTR (Supplementary Fig. S5G, H). Together, these data revealed that NXPH1/α-NRXN1 signaling is necessary and sufficient for NB tumor growth *in vivo* (Fig. 4K).

### NXPH1/α-NRXN signaling inhibits the metastatic potential of NBs

Finally, we set out to define whether NXPH1/α-NRXN signaling regulates the metastatic potential of NB cells *in vivo*. To this end, we generated clones of SK-N-SH cells carrying a constitutively-expressed luciferase cassette in addition to the doxycycline-inducible sh-RNAs and eGFP. FACS-purified, doxycycline-treated cells from the sh-NXPH1, sh-αNRXN1 and sh-Ctrl clones were injected into the left cardiac ventricle of immunodeficient NOD-SCID gamma (NSG) mice and their dissemination and metastatic growth were monitored non-invasively by bioluminescence (BLI) over 9 weeks (Fig. 5A). A few minutes after injection, cells from the sh-Ctrl, sh-NXPH1 and sh-αNRXN1 clones were seen disseminating within the body of the mice (Fig. 5B). Their BLI then regressed to background levels during the first 3-4 weeks, indicating that the vast majority of the cells disappeared during that period (Fig. 5B). After 9 weeks, 86% and 57% of the mice injected with sh-NXPH1 and sh-αNRXN1 cells had developed detectable metastatic tumor masses, whereas only 29% of the mice injected with the sh-Ctrl cells had (Fig. 5B, C). Moreover, the metastatic tumors developed by the sh-NXPH1 and sh-αNRXN1 cells were detected earlier and produced a photon count much higher than the sh-Ctrl cells (Fig. 5B, D). Metastases were particularly evident in the liver and bone marrow, in agreement with previous reports on the organotropism of metastatic NBs (5, 45). Thus, reducing the expression of NXPH1 or that of its α-NRXN1 receptor stimulated the metastatic potential of NB cells *in vivo*. These latter findings therefore revealed that NXPH1/α-NRXN1 signaling represses the dissemination and metastatic potential of NBs (Fig. 5E).

**Figure 5:**
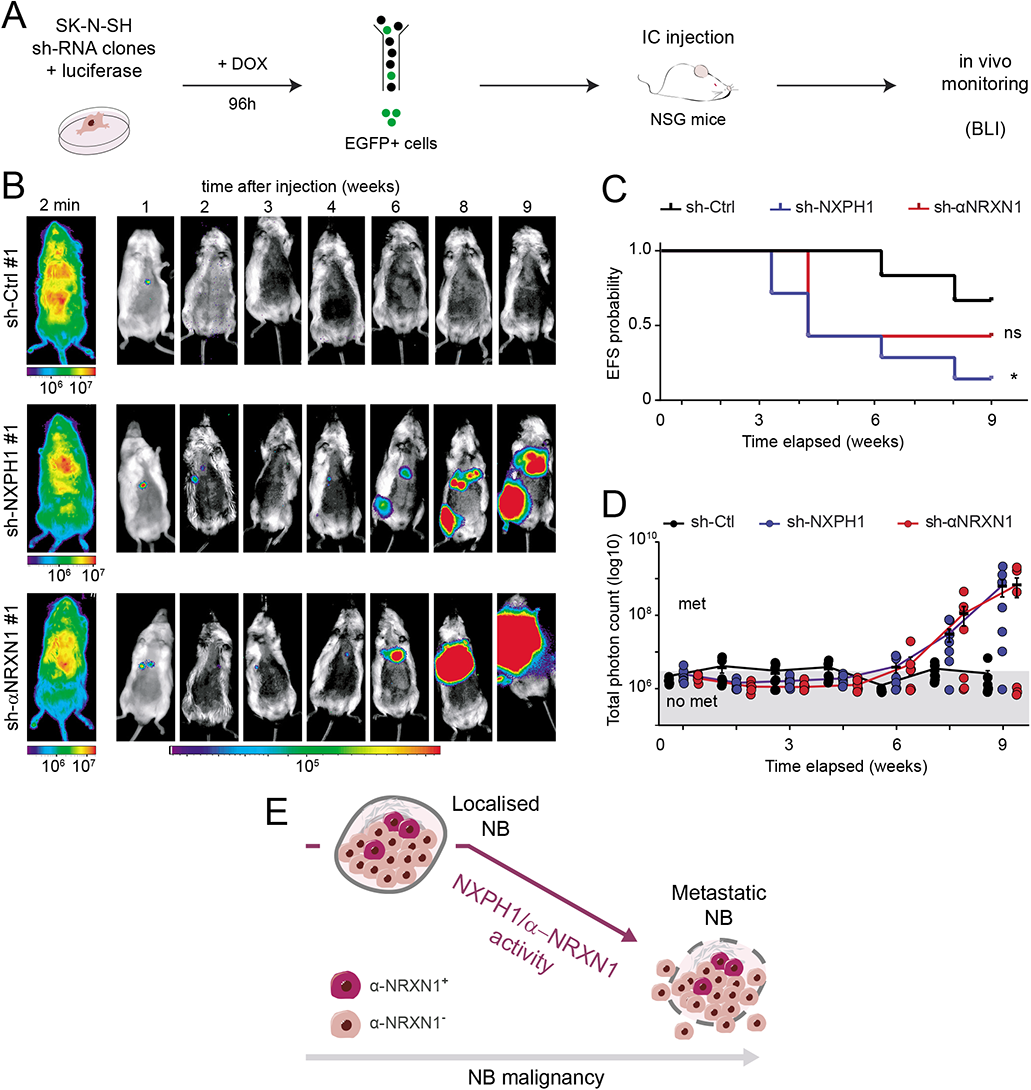
The inhibition of NXPH1 or α-NRXN1 activity unleashes the metastatic potential of NB cells. (**A**) The capacity of pre-activated (treated with doxycyline for 96 hours *in vitro*, + DOX) and FACS-purified sh-Ctrl, sh-NXPH1 and sh-αNRXN1 cells to develop metastases was assessed by luciferase-derived bioluminescence after intra-cardiac injection in immunodeficient NSG mice. (**B**) Representative images of the *in vivo* luciferase-derived bioluminescence detected in animals 2 minutes or 1 to 9 weeks after injection with sh-Ctrl, sh-NXPH1 or sh-αNRXN1 cells. (**C, D**) Follow-up of (**C**) the EFS and (**D**) the total photon count (± s.e.m) quantified in NSG mice injected with sh-Ctrl (black, n=7), sh-NXPH1 (blue, n=7) or sh-αNRXN1 (red, n=7) cells. Each dot represents the photon count of a single animal. The grey area defines the background bioluminescence. (**E**) Model proposing that the downregulation of NXPH1/α-NRXN1 activity in NB tumors facilitates their metastatic potential and dissemination. Statistical significance was assessed using a non-parametric Mann-Whitney test (**C**) or a Chi-squared test (**D**): *p<0.05; ns, p>0.05.

## Discussion

Our study provides fundamental insights into the transcriptional signatures associated with both the aetiology and malignancy of NB. We notably reveal that a signature corresponding to the core SA lineage strongly distinguishes LR-NBs from HR-NBs, this signature being largely composed of genes associated with favorable patient outcome and with reduced disease progression. Encoded by one of these SA-c1 genes, the secreted protein NXPH1 and its receptor α-NRXN1 stimulate the growth of NB cells but markedly restrict their metastatic potential.

The pathogenesis of NB remains puzzling for several reasons. First, some NB tumors can regress spontaneously (46, 47). Second, so far there is little evidence showing that LR-NBs ever evolve into HR-NBs once diagnosed (48). This led to the idea that LR-NBs and HR-NBs represent distinct types of tumors and that they likely originate at different stages of development within the neural crest lineage. Various studies recently employed single-cell transcriptomics to tackle this question and to identify the NB cell-of-origin, which yielded discrepant conclusions (12–16,49–52). Whereas one study argued that adrenal NBs derive from fetal chromaffin cells and that LR-NBs and HR-NBs can be discriminated based on the degree of chromaffin cell differentation (12), others pinpointed instead fetal sympathoblasts as the NB cell-of-origin (11,14–16). Another study found that the signatures of sympathoblasts and norepinephrine-chromaffin cells were highly enriched in samples from NB patients, but suggested that the signature of Schwann cell precursors is the one distinguishing LR-NBs from HR-NBs (13). As we herein focused directly on identifying the transcriptional signature associated to NB aetiology that further distinguishes LR-NBs and HR-NBs, we did not attempt to settle this current debate. We specifically addressed the component of the SA lineage that distinguishes LR-NBs and HR-NBs, using transcriptional signatures for human SCPs, bridge cells, chromaffin cells and sympathoblasts recently identified by single-cell RNA-seq (16). In contrast to previous reports (12–16), we tested these signatures on a restricted list of LR-NB *vs* HR-NBs DEGs and observed that these 4 signatures all showed a strong enrichment for genes expressed at higher levels in LR-NBs than in HR-NBs. More importantly, having realised that the SCP, bridge cell, chromaffin cell and sympathoblast signatures that we used were largely overlapping, we then demonstrated that their enrichment for genes expressed at higher levels in LR-NBs is due to the signature shared by these sympatho-adrenal cell identities, rather than to the transcriptional singularities of any of these four cell types. Notably, this core SA signature retrieved more than half of the 503 LR *vs* HR DEGs, highlighting the importance of this developmental signature in discriminating the transcriptional landscapes of LR-NB and HR-NBs. Remarkably, this core SA signature retrieved 72% of the LR *vs* HR DEGs forming the cluster c1 (i.e. the genes expressed at higher levels in LR-NBs than in HR-NBs than in fAG samples and associated with a favorable patient outcome). This preferential correlation between the transcriptional identity of the SA lineage and the LR-NB phenotype suggests that this developmental program actually opposes NB malignancy, which was unforeseen. Put together, our findings shed a new light on the complex contribution of the neural crest-derived developmental program to NB pathogenesis. The SA program indeed appears to facilitate NB formation but to concomitantly restrict its malignant potential. The dual impact of NXPH1/α-NRXN signalling on NB cell behaviour presented here is a first experimental illustration of this unexpected model.

*NXPH1* came out in our genome-wide transcriptomic analysis as the SA-c1 gene with the second highest enrichment in LR-NBs relative to HR-NBs. The intensity of *NXPH1* expression stratifies NB patients relative to their tumor stage and to the probability of disease progression, extending findings reported in a genome-wide associated study that specifically compared INSS-3 and INSS-4 NB tumors (53). Interestingly, *NXPH1* can also be used as a DNA methylation biomarker associated with a good prognosis for NB patients (54), and represents one of the genes differentially expressed between LR-NBs and HR-NBs with the most restricted expression pattern in postnatal human tissues (16). Therefore, monitoring *NXPH1* levels might be clinically relevant for the therapeutic management of patients suffering from NB.

The secreted glycoprotein NXPH1 specifically binds and modulates the activity of the transmembrane receptors α-NRXNs (34–37). The fact that *NRXN2* also came out as a SA-c1 gene and that the expression levels of both *NRXN1* and *NRXN2* correlate to a better prognosis is noteworthy. Although the data available did not allow us to determine if these correlations were directly related to *α-NRXN* isoforms, these findings motivated our choice to focus on NXPH1/α-NRXN signaling for functional studies. The results from our functional assays demonstrated that both NXPH1 and α-NRXN1 regulate various aspects of NB cell behaviour. First, they stimulate NB tumor growth and we provide evidence that NXPH1 fuels the proliferation of NB cells. Second, NXPH1 and α-NRXN1 knockdowns both unleash the ability of NB cells to colonise tissues like the liver and bone marrow, two of the main metastatic organs in HR-NB patients (5, 45). The molecular mechanisms causing these two opposite effects remain unclear and their study is made harder by the fact that the signaling cascade triggered downstream of α-NRXNs in response to NXPH1 binding is as yet unknown. Nevertheless, this unexpected anti-metastatic activity of NXPH1/α-NRXN signaling likely explains why higher *NXPH1* and α*NRXN1/2* expression levels are correlated with a better patient prognosis. Targeting and modulating NXPH1/α-NRXN signaling might thus represent an appealing strategy to treat metastatic, therapy-resistant HR-NBs.

In summary, our work reinforces the importance of the neural crest-derived SA developmental program in NB pathogenesis and especially highlights the necessity to examine how these genes regulate both the growth and the metastatic potential of NBs. This will facilitate the identification of novel actors, such as NXPH1/α-NRXN1 signaling, that might serve as therapeutic targets to treat children affected by metastatic, therapy-resistant HR-NBs.

## Materials and Methods

### Animal Studies

Fertilized white Leghorn chicken eggs were provided by Granja Gibert, rambla Regueral, S/N, 43850 Cambrils, Spain. Eggs were incubated in a humidified atmosphere at 38°C in a Javier Masalles 240N incubator, manipulated at embryonic day 10 (E10) and sacrificed at E17. Sex was not identified.

5 weeks-old female NOD/SCID mice (NOD.CB17-Prkdc^scid^ /NCrHsd; RRID: IMSR_ARC:NODSCID) and 7 weeks-old male NSG mice (NOD.Cg-provided by Envigo were used for subcutaneous xenografting and metastasis analyses, respectively. Mice were housed under a regimen of 12 hours light/12 hours dark cycles in specific pathogen-free conditions. Sterile dry pellets and water was administered *ad libitum.* Animals were sacrificed by cervical dislocation or CO_2_ administration at the end of the experiment or when required for ethical reasons.

#### Patient-derived xenografts

Human NB-PDX models (HSJD-NB-012, HSJD-NB-011 and HSJD-NB-007) were established, maintained and provided by A.M.C under a local animal care and use committee-approved protocol (135/11). All NB-PDX models were derived from patients with progressive stage INSS4 disease, refractory to all treatments. HSJD-NB-011 was established from a 2.5 year old male, with amplification of MYCN gene, TP53 wild type and ALK-mutated (I1171N). HSJD-NB-007 was established from a 5 year old male with amplification of MYCN gene and no mutations in TP53 and ALK. HSJD-NB-012 was established from a 4 year old male and its genetic profile is unknown. HSJD-NB-011 and HSJD-NB-007 have been previously used for publication (18, 19). Experimental manipulation of NB-PDXs was performed in HSJD under the supervision of A.M.C.

To quantify α-NRXN1 expression in PDXs, tumours were dissected off the flank of immunocompromised nude mice, minced using razor blades in a sterile dish and transferred to a 50ml falcon tube. Cells were dissociated following manufacturer instructions using the Brain Tumour Dissociation Kit and a gentle MACS dissociator (Milteny Biotec #130-095-942 and #130-093-235). Red blood cells were removed by using ACK buffer. A single cell suspension weas prepared in FACS buffer (ice-cold DMEM-5%FBS) containing 10μM of Rock inhibitor (SIGMA #Y0503) to a final concentration of 1·10^7^ cells/ml and stained for flow cytometry analysis as explained below

#### Cell lines

HEK-293 (RRID: CVCL_0045), LAN-1 (RRID: CVCL_1827), IMR-5 (RRID: CVCL_1306), SH-SY5Y (RRID: CVCL_0019), SK-N-AS (RRID: CVCL_1700), SK-N-JD (RRID: CVCL_WH12), SK-N-LP (RRID: CVCL_WH13), LA1-5s (RRID: CVCL_2549), SK-N-SH (RRID: CVCL_0531), SL-N-Be(2)c (RRID: CVCL_0529) and IMR-32 (RRID: CVCL_0346) cell lines were obtained from Marian Martínez-Balbás (IBMB-CSIC), Cinzia Lavarino (HSJD) and Joan Xavier Comella (VHIR). In basal culture conditions, NB cell lines were grown in either RPMI 1640-Glutamax (ThermoFisher Scientific cat# 61870010) supplemented with 10% (LAN-1, IMR-5, SH-SY5Y, SK-N-AS, SK-N-JD, SK-N-LP, LA1-5s) or 20% (SK-N-SH, SK-N-Be(2)c, IMR-32) foetal bovine serum (FBS; Cultek cat# 16SV30160.03RYB35908) plus 1% penicillin/streptomycin antibiotic cocktail (ThermoFisher Scientific cat# 15140-122). The HEK-293 cell line was grown in DMEM-Glutamax (Lifetechnologies cat#61965-026) supplemented 10% FBS and 1% penicillin/streptomycin cocktail. Unless indicated, cells were grown in standard culture conditions. The presence of mycoplasma contamination was routinely checked by immunofluorescence. Cell lines have not been re-authenticated for the present paper.

#### Genome-wide transcriptomic analysis

The HSJD-NB dataset used in this study has been published previously (20) and is available at GEO repository (GSE54720). The full human SCP, bridge cell, chromaffin cell and sympathoblast signatures were obtained from a published study (16), and the complementary signatures were obtained from a Venn diagram analysis performed with a freely available software developed by the group of Dr Y. Van de Peer (VIB, Brussels, Belgium, http://bioinformatics.psb.ugent.be/webtools/Venn/).

Genome-wide transcriptomic analyses were performed using the web-based Phantasus software (https://artyomovlab.wustl.edu/phantasus; version 1.5.1) (21). Normalized log expression values were obtained using the Quantile Normalize Adjustment tool. The Maximun Median Probe method was then selected to collapse the different probes of a single gene. Finally, genes were filtered based on raw expression levels and the 12,000 most-expressed genes from each array were selected for subsequent analysis. Sample dispersion was then assessed using a principal component analysis leading to the removal of one outlier (#LR-08) from further analysis. Differentially-expressed genes (DEGs) between groups were identified using the Limma R package. The genes were considered as differentially expressed when the false discovery rate (FDR)-adjusted P-value (with Benjamini-Hochberg procedure) was <0.05. Gene set enrichment analyses (GSEA) were performed using the FGSEA tool from Phantasus.

#### Gene ontology analysis

Gene ontology (GO) term enrichment analyses (biological process) were performed with the PANTHER classification system (http://pantherdb.org) (22, 23). The 50 most-enriched GO terms were used to assess GO term distribution and were annotated with the Ancestor Chart tool of QuickGO (https://www.ebi.ac.uk/QuickGO).

#### R2 genomic analysis and visualization platform

Publically available NB patient microarray SEQC 498 (GSE62564) was obtained from the R2 genome analysis and visualization platform (https://hgserver1.amc.nl/cgi-bin/r2/main.cgi) (24). The R2-web based application was used to generate Kaplan-Meier event-free survival curves. Data were grouped based on the expression levels of either *NXPH1, NRXN1-3* or the full SA-c1 gene module. In the latter case, the SEQC cohort was subdivided into quartiles based on the combined expression of the 242 SA-c1 genes, considering as “high” or “low” the expression of each SA-c1 gene with respect to its own average expression level. The quartiles Q1-Q4 thereby consisted of 123-126 samples showing numbers of SA-c1 genes expressed at high levels as follows: 487≤Q1≤360; 359≤Q2≤250; 249≤Q3≤135 and 134≤Q4≤4. Statistical significance was automatically assessed by the R2-server using a log-rank test. The R2-web based application was also used to compare the distribution of patients among different NB prognosis groups according to expression levels of the SA-c1 gene module or *NXPH1.* For the analysis of disease progression, the category “progression” included tumors that did not respond to therapy plus those that were recurrent despite an initial therapeutic response. Statistical significance was automatically assessed by R2-server using the Chi-square + Fisher’s test.

#### Plasmids and lentiviral infection

Inhibition of *NXPH1* and *α-NRXN1* expression was triggered by lentiviral infection of short-hairpin constructs inserted into in pLKO.1 or doxycycline-inducible pSLIK-Neo vectors (25, 26). The pLKO.1-TRC cloning vector (RRID: Addgene_10878) and the pLKO.1-sh-Ctl (RRID: Addgene_10879) were kindly provided by Marian Martínez-Balbás. Doxycycline-inducible miRshRNA-eGFP-expressing pSLIK-Neo lentiviral vectors were generated by gateway recombination between the TTRE-eGFP-miR-shRNA entry vector and the pSLIK destination vector (26). pEN_TTGmiRc2 and pSLIK-Neo were purchased from Addgene (Cat#25753, RRID: Addgene_25752 and Cat#25735, RRID: Addgene_25735; gifts from Ian Fraser). *NXPH1* and *α-NRXN1-* specific sh-RNA sequences were designed using the web-based Genetic Perturbation Platform of the Broad Institute (https://portals.broadinstitute.org/gpp/public/). Expression of luciferase was triggered by lentiviral infection of pLEX-hFL2iG vector.

Lentiviral particles were generated using the psPAX2 (RRID: Addgene_12260) and pMD2.G plasmids (RRID: Addgene_12259) kindly provided by Marian Martínez-Balbás (IBMB-CSIC). Briefly, HEK293T cell line was transfected using the Lipofectamine 2000 Tranfection Reagent (ThermoFisher Scientific #11668019) following manufacturer’s instruction and lentiviruses were harvested 48h post-transfection. Transduction of target cells was usually performed readily after lentivirus recovery. 2μg/ml of puromycin (SIGMA #p8833) and 600μg/ml of neomycin/G418 (SIGMA-ALDRICH #A1720) were used for selection of transduced cells.

#### RNA purification and RT-qPCR

Total RNA was extracted with TRIzol Reagent (ThermoFisher Scientific #15596-018) and Pellet Paint Co-precipitant (Merck Millipore #69049-3). Genomic DNA traces were removed with DNA-free DNase Treatment and Removal Reagents (FisherScientific #AM1906) following manufacturer’s instructions. DNA-free RNA was retro-transcribed using the High-Capacity cDNA Reverse Transcription Kit (LifeTechnlogies #4368814). Semi-quantitative PCR was performed using the Lightcycler 480 SYBR Green 2x Master (Roche #04887352001) on a Roche Lightcycler 480 Real-Time PCR System. Unless indicated otherwise, primers were design using the UCSC Genome browser and Primer3Plus online softwares. *18SrRNA* or *TBP* were used as housekeeping genes. Relative cDNA levels were calculated using the efficiency-corrected ΔCt method (27). Heatmap representations were performed using GraphPad Prism v6 (RRID: SCR_002798).

#### Fluorescence-activated cell sorting (FACS)

All experiments were conducted using a BD FACSAria Fusion at the Cytometry Core Facility (Centres Científics i Tecnològics) of Universitat de Barcelona. Briefly, single cell suspensions coming from *in vitro* cell cultures or dissociated PDXs were prepared in FACS buffer (ice-cold DMEM-5%FBS) at 10^7^ cells/ml and incubated for 60min at 4°C with the following fluorescent-conjugated antibodies diluted in FACS buffer: rat monoclonal anti-CD31 (clone 390), PE-Cy™7 (BD Pharmingen Cat#561410, RRID: AB_10612003), rat monoclonal anti-CD11b (clone M1/70), PE-Cy™7 (BD PharMingen Cat#561098, RRID: AB_2033994), mouse monoclonal anti-Disialoganglioside GD2 (clone 14.G2a), Alexa Fluor® 647 (BD PharMingen Cat#562096, RRID: AB_1154051) and a rabbit polyclonal anti-Neurexin 1α, ATTO488 (Alomone Labs Cat#ANR-031-AG, RRID: AB_2756687). Finally, cells were washed twice with ice-cold FACS washing buffer and resuspended at 5·10^6^ cells/ml in ice-cold FACS buffer. Dox-induced cells from sh-Ctrl, sh-NXPH1 and sh-αNRXN1 clones were purified based on eGFP sorting. DAPI (0.1μg/ml) or propidium iodide (2μg/ml) were used as vital dyes

The gating strategy for detection and purification of α-NRXN1 subpopulations was adapted from a previous study (28). The 10-12% of NB cells showing the strongest mean fluorescent intensity were used as α-NRXN1^+^ and the 20-40% with the lowest intensity were used as α-NRXN1^-^. Among α-NRXN1^+^ cells, the 0.5-1% of cells showing the highest intensity were considered as α-NRXN1^High^ and the 10-20% with an intensity 3 to 4 times lower as α-NRXN1^Low^. For depletion experiments, the 10-15% of cells showing the brightest NRXN1-A488 intensity was discarded. FACS data was analyzed with FlowJo (RRID: SCR_000410).

#### MTT assay

Approximately 8,000 cells were seeded in quadruplicates. At desired timepoints, 10μl of 5mg/ml MTT solution (SIGMA #M2128) was added to each well and incubated at 37°C for 4 hours. Formazan crystals were dissolved with 100μl DMSO (SIGMA #D2650) and the optical density at 570nm (OD570) was measured. This assay was performed on n=2-4 biological replicates.

#### S-phase index and BrdU-retention assay

A solution containing 10 μM of 5-bromo-2’-deoxyuridine (BrdU; SIGMA #858811) was added to the growth medium of SK-N-SH cell monolayers and cells were incubated at 37°C for 2 hours to estimate the S-phase index, or for 24 hours prior to the CAM assay to assess BrdU retention, using n=4-9 biological replicates. Cells were then fixed, permebilized and incubated with 5U/ml DNase I type II (SIGMA #D4527) for 15 min at RT. BrdU incorporation was then assessed by immunofluorescence as described below.

#### Tumour spheroid assay

To test the sphere-forming capacity of the different human NB cell lines, cells were grown in restrictive culture conditions using a media containing DMEM/F12 (ThermoFisher Scientific #11330-032), 10% KnockOut serum replacement (ThermoFisher Scientific #10828012), 2mM Glutamax (ThermoFisher Scientific #35050038), 1% MEM non-essential amino acids (ThermoFisher Scientific #11140035), 0.1mM β-mercaptoethanol (SIGMA-ALDRICH #M3148), 1% penicillin/streptomycin cocktail (ThermoFisher Scientific #15140-122) and 0.1ng/ml basic FGF (ThermoFisher Scientific #PHG0021). Medium was renewed every week. The sphere-forming capacity of the different human NB cell lines was performed on n=2 biological replicates.

To assess mRNA expression in stem cell-enrichment conditions, 3·10^6^ cells were grown on uncoated 60mm plates in 2ml SFM. Spheres were allowed to grow for 5 weeks and were photographed using an inverted bright-field microscope (Leica DMIRBE). Viable spheres were recovered by centrifugation at 100g for 5min at 4°C) and processed for RNA extraction as described below.

To assess the sphere-forming capacity of NB cells, 2,000 dissociated cells from each population of interest were sorted in quadruplicates and seeded directly onto 12-well plates coated with 0.5% agarose (Condisa Laboratory #8014). Cells were allowed to grow for 1 week and the presence of spheroids was monitored every two days. To detect the presence of viable spheroids after 1 week, plates were incubated with 0.5mg/ml MTT (SIGMA #M2128) for 3 hours at 37°C. Viable spheres (black coloured) were manually counted using an inverted bright-field microscope (Leica DMIRBE). The sphere-forming capacity was assessed for n=1-4 biological replicates per experimental condition.

#### 2D Extreme limiting dilution assay (ELDA)

Cells subpopulations were FACS-sorted, seeded in triplicates into 96-well plates and grown until one experimental condition fully covered the well surface. Half of the growth medium was renewed every week. Cells were then fixed with 4% PFA (SIGMA #16005) for 15min at RT, washed with PBS and stained with 0.5% crystal violet (SIGMA #V5365) for 20 min at RT. The dye excess was washed out and plates were air-dried for at least 2 hours at RT. Crystal violet staining was dissolved by adding 200μl methanol to each well, followed by 20min at RT. Finally, the OD570 was measured. This assay was performed on n=3 biological replicates per experimental condition.

#### Immunofluorescence

Fixed samples (either cells growing *in vitro* or tumor sections) were permeabilized with 0.1% Triton X100 in PBS (0.1% PBT) for 15 minutes at RT, saturated for 30 min in a blocking buffer (10% horse serum, 1% BSA, 0,3% Glycine in 0.1% PBT), and incubated overnight at 4°C with primary antibodies diluted in blocking buffer. After 3 washes in PBS, samples were incubated with fluorescence-conjugated secondary antibodies for 2 hours at RT. Samples were stained with rat monoclonal anti-BrdU [clone BU1/75 (ICR1)] (BioRad Cat #MCA 2060, RRID: AB_323427), rabbit monoclonal anti-active Caspase-3 (clone C92-605) (BD PharMingen Cat#559596, RRID: AB_397274), rabbit polyclonal anti-phospho-Histone H3 (Ser10) (Upstate Cat#06-570, RRID: AB_310177), rat monoclonal anti-phospho-Histone H3 (pSer28) (clone HTA28) (Sigma Cat#H9908, RRID: AB_260096), mouse monoclonal anti-Human Nuclear Antigen (clone 235-1) (Abcam Cat#ab191181, RRID: AB_2492189), rabbit polyclonal anti-Ki67 (Abcam Cat#ab66155; RRID: AB:1140752), mouse monoclonal anti-p75NTR (clone ME20.4) (Millipore Cat#05-446, RRID: AB_309737) and mouse monoclonal anti-PSA-NCAM (clone 2-2B) (AbCys Cat#AbC0019, RRID: AB_2313692). Cell nuclei were stained with 1 μg/ml DAPI and samples mounted in home-made Mowiol. At least n=4 independent samples were assessed for each experimental condition.

#### Image acquisition and treatment

Optical sections of human NB cell samples were acquired at RT with the Leica LAS_X (2016) software on an Automated Inverted Leica AF7000 wide-field microscope using 20x (Dry/NA 0.5/HC PL APO/0.70 CS∞/0.17/C) or 40x (oil/NA 1.25-0.75/HCX PL APO Lbd Blue ∞/0.17/D). Cell counting was performed in minimum of 4 fields per well using the ImageJ software (https://imagej.nih.gov/ij/;RRID: SCR_003070) (29, 30).

Optical sections of NB tumour graft samples were acquired at RT with the Leica LAS software (RRID: SCR_013673), in a Leica TCS SP5 confocal microscope using 20x (dry/HC PL APO 20x/0.70 CS ∞/0.17/C) or 40x (oil /HCX PL APO 40X/1.25-0.75 OIL CS ∞/0.17/D) objective lenses or with the LSM Software ZEN 2.1 (RRID: SCR_018163), in a ZEISS Lsm780 confocal microscope using 25x (oil,w,Glic, NA 0.8, Plan-Apochromat/ImmKorr) or 40x (oil/NA 1.3/Plan-Apochromat/(UV)VIS-IR). Whole sections of NB tumour grafts were acquired as tile scans using the motorized XY stage. Tiles were directly stitched into a single mosaic by the microscope software. Maximal projections obtained from 3-6μm Z-stack images were processed with ImageJ for image merging, resizing and cell counting. Cell counting was performed in a minimum of two sections per tumour.

#### Cell counting

Two Fiji macros were developed for cell counting. The first one uses the DAPI channel to create a binary mask containing all the nuclei; segmentation is based on local contrast enhancing, filtering and a manual pipeline for fine tuning. The second one uses the so-produced binary mask to extract the nuclei selections and collate them against the original images, in order to intensity threshold the nuclei for the three markers used: GFP, Ki67 and pH3. The macro delivers a result table and a processed compound image with all the nuclei labelled according to their content. Codes and the details on the procedure can be downloaded at: https://github.com/MolecularImagingPlatformIBMB/CellProliferationAssay.

#### Tumour Xenografts

SK-N-SH clones carrying a conditional knockdown of *NXPH1 (*sh-NXPH1), α-NRXN1 (sh-αNRXN1) or a scramble control sequence (sh-Ctrl) were first activated *in vitro* in presence of 1mg/ml doxycycline (ACEFESA #PAA29510025) for 96 hours. Pre-activated cells were then sorted by FACS based on eGFP expression, and resuspended in grafting media consisting of DMEM/F12, 7mg/ml Matrigel (Fisher Scientific #10365602, 1mg/ml doxycycline and 10μM of Rock inhibitor (SIGMA #Y0503) at a concentration of 0.5·10^7^ cells/ml. Then, 100μl cell suspensions were grafted subcutaneously using a 0.5ml BD Micro-Fine 29G insulin syringe. Each animal received 2 grafts, one on each flanks.. A fresh solution containing 2mg/ml of doxycycline dissolved in 7.5% sucrose-bearing sterile water was administered *ad libitum* in drinking water, its administration starting 2 days prior to xenografting. Mouse weight and tumour growth were monitored twice a week until tumour detection. From this point, the tumour volume was measured every two days with a digital calliper and mice were sacrificed once the tumour volume reached ethical permission limits. Tumours were then resected, weighted and photographed.

#### Experimental metastasis assay

SK-N-SH clones carrying a luciferase cassette and a conditional knockdown of *NXPH1 (*sh-NXPH1), α-NRXN1 (sh-αNRXN1) or a scramble control sequence (sh-Ctrl) were first activated *in vitro* in presence of 1mg/ml doxycycline for 96 hours. To test the metastatic potential of NB cells, 2·10^5^ cells resuspended in 100µl of PBS were injected into the left cardiac ventricle of 7 weeks-old male NSG mice (NOD.Cg-Prkdcscid Il2rgtm1Wjl/SzJ) with a 26G ½ 13mm needle, using a total of 7 injected mice per experimental condition. 2mg/ml of doxycycline dissolved in 7.5% sucrose-bearing sterile water was administered *ad libitum* in drinking water as explained before. The generation of metastases was monitored twice per week by bioluminescence imaging (BLI) with the IVIS Imaging System (Perkin Elmer) after retro-orbital administration of 15µg/µl D-Luciferin (GoldBio #LUCK-10G). Metastatic lesions were quantified by measuring the photon count of the region of interest, using an exposure time of 2 minutes and a medium binning value. Images taken at time “2 min” post-injection are presented with a higher threshold (10^7) than images taken at later time points to show the even cell distribution throughout the body right after injection.

#### Chick CAM assay

At E10, a small window was created on the egg shell using a sterile scalpel. Cells were resuspended in grafting media consisting of DMEM/F12, 7mg/ml Matrigel and 10μM of Rock inhibitor were implanted onto the surface of the chorio-allantoid membrane (CAM) by gently scraping the upper CAM layer (avoiding bleeding or visible rupture of the capillaries). To assess the effects of NXPH1 on tumour growth, parental SK-N-SH cells were trypsinized and resuspended in grafting media in presence of 10μg/ml recombinant NXPH1 (rNXPH1) or 100μg/ml BSA (Ctrl; SIGMA #A7906). To test the growth of NB cells depleted from their α-NRXN1^+^ subpopulation, parental SK-N-SH depleted or not from their α-NRXN1^+^ subpopulation were FACS-purified and resuspended in grafting media. In both experiments, a concentration of 0.5·10^6^ cells/10μl was used and grafted onto the CAM, using n=4-13 biological replicates. At E17 NB tumour grafts were removed from the CAM and tumours were photographed, weighed and measured using a digital calliper (VWR). The final tumour volume (V) was calculated using the following formula: V= 4/3·π·length·depth·width. Tumours were then fixed in 4% PFA for 1h30min at 4°C, washed in PBS1x and sequentially cryo-protected with PBS solutions containing 15% and 30% sucrose. Small cubical tumour pieces (5mm·5mm·5 mm approx.) were then embedded in Tissue-tek (Sakura Finetek #4583) and frozen on dry ice. Tumour pieces were sectioned on a cryostat (Leica) at 16μm, collected serially on home-made TESPA pre-coated slides and slides stored at −20°C. Immunofluorescence was performed as described above.

#### Statistics

No statistical method was used to predetermine sample size. The experiments were not randomized. The investigators were not blinded to allocation during experiments. Statistical analyses were performed using the GraphPad Prism 6 software (RRID: SCR_002798). Unless noted otherwise (see quantifications), cell counts were typically performed on 3 images per sample and *n* values correspond to different tumors or biological replicates. The normal distribution of the values was assessed by the Shapiro-Wilk normality test. Significance was then assessed with a two-way ANOVA + Dunnett’s or Tukey’s test for data presenting a normal distribution, or alternatively with the non-parametric Mann–Whitney test, the Mantel-Cox log-rank test or the Kruskal-Wallis test + Dunn’s test for non-normally distributed data. For survival analysis, significance was assessed with the Gehan-Breslow-Wilcoxon test. Statistical analysis performed on the GSE62564 dataset was automatically assessed by the R2 server (https://hgserver1.amc.nl/cgi-bin/r2/main.cgi) using the Mantel-Cox log-rank test or the Chi-square + Fisher’s test. The following convention was used: n.s: P>0.05; *P<0.05, **P<0.01, ***P<0.001. Statistical details of experiments can be found in the Fig. legends and main text above.

## Acknowledgements

**Ethics approval and consent to participate:** All the experimental procedures involving mice were carried out in accordance with the European Union guidelines (Directive 2010/63/EU) and according to the guidelines from the Animal Care Committee at the Generalitat de Catalunya. The studies were approved by the ethics committee of the Parc Científic de Barcelona and by the Parc de Recerca Biomèdica de Barcelona (PRBB) animal facility policy. According to Spanish animal care guidelines, no approval was required to perform the experiments involving chicken embryos herein. Human NB-PDX models (HSJD-NB-012, HSJD-NB-011 and HSJD-NB-007) were established, maintained and provided by A.M.C under a local animal care and use committee-approved protocol (135/11).

**Acknowledgements:** we thank the members of the laboratories of E.M., Maria L. Arbonés and Sebastian Pons (IBMB-CSIC) for discussions related to this study. We thank the Xarxa de Bancs de Tumors de Catalunya (XBTC; sponsored by Pla Director d’Oncologia de Catalunya), the “Biobanc de l’Hospital Infantil Sant Joan de Déu per a la Investigació” integrated in the National Network Biobanks of ISCIII for the sample and data procurement, the IBMB Molecular Imaging platform and the PCB Flow Cytometry facility for their assistance. We are also grateful to Marian Martínez-Balbás (IBMB-CSIC) and Joan Xavier Comella (Vall d’Hebron Institut de Recerca, Barcelona, Spain) for providing reagents.

## Author contributions

**Conceptualization:** L.F-E. and G.L.D.; **Methodology:** L.F-E., I.S., S.G-G., E.L.G, S.U., E.R., M.D.M. and G.L.D.; **Investigation:** L.F-E., I.S., S.G-G., E.L.G and G.L.D.; **Resources:** A.M.C., T.C-T., C.L. and E.M; **Visualization:** L.F-E. and G.L.D.; **Writing - Original Draft:** L.F-E. and G.L.D.; **Funding Acquisition:** E.L.G., A.M.C., T.C-T., C.L. and E.M. **Supervision:** E.M. and G.L.D.

## Availability of data and materials

All data generated or analysed during this study are included in this article and its supplementary information files.

## Competing interests

The authors have no competing financial interests to declare.

**Correspondence and requests for materials** should be addressed to G.L.D.

## Supplementary files

**Supplemental Figure S1:**
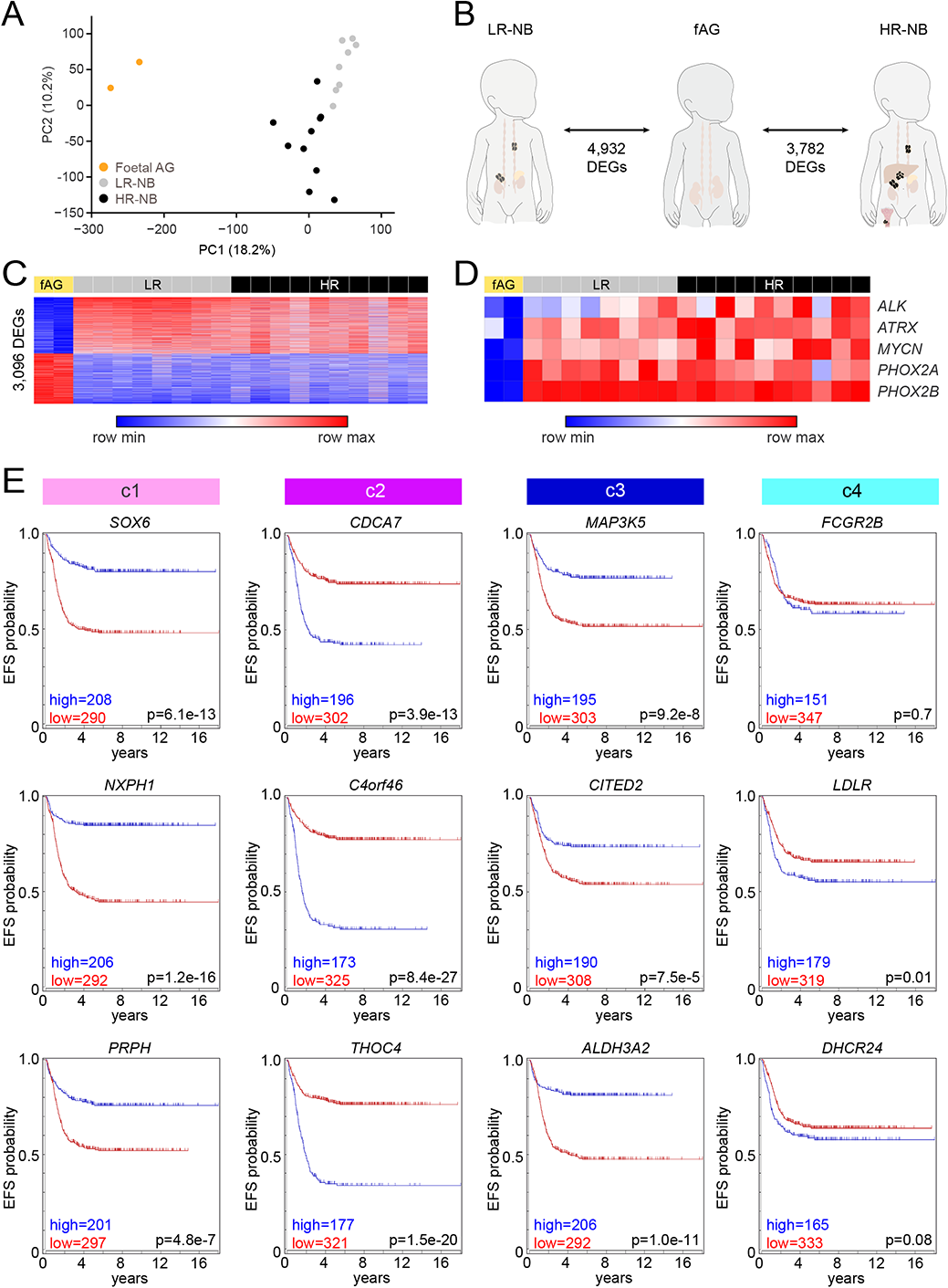
Genome-wide transcriptomic analysis of neuroblastoma formation and malignancy and event-free survival probability of the top 3 candidates genes of clusters c1-c4. (A) Principal component analysis performed on our panel of LR-NBs, HR-NBs and human fetal adrenal gland samples. (B) Numbers of DEGs retrieved after comparing LR vs fAG and fAG vs HR. (C, D) Heatmaps of (C) the total list of DEGs common to the independent comparisons LR vs fAG and fAG vs HR (adj. p<0.05) and (D) known markers of NB malignancy (MYCN, ALK, ATRX, PHOX2A and PHOX2B). (E) Follow-up of the event-free survival (EFS) probability observed for a cohort of 498 NB patients (SEQC database, R2 online platform) subdivided based on the expression levels of the top-3 differentially expressed genes from each cluster identified (c1-c4). The statistical significance was assessed automatically by the R2 server using a log-rank test. The exact p-values are indicated in each graph.

**Supplemental Figure S2:**
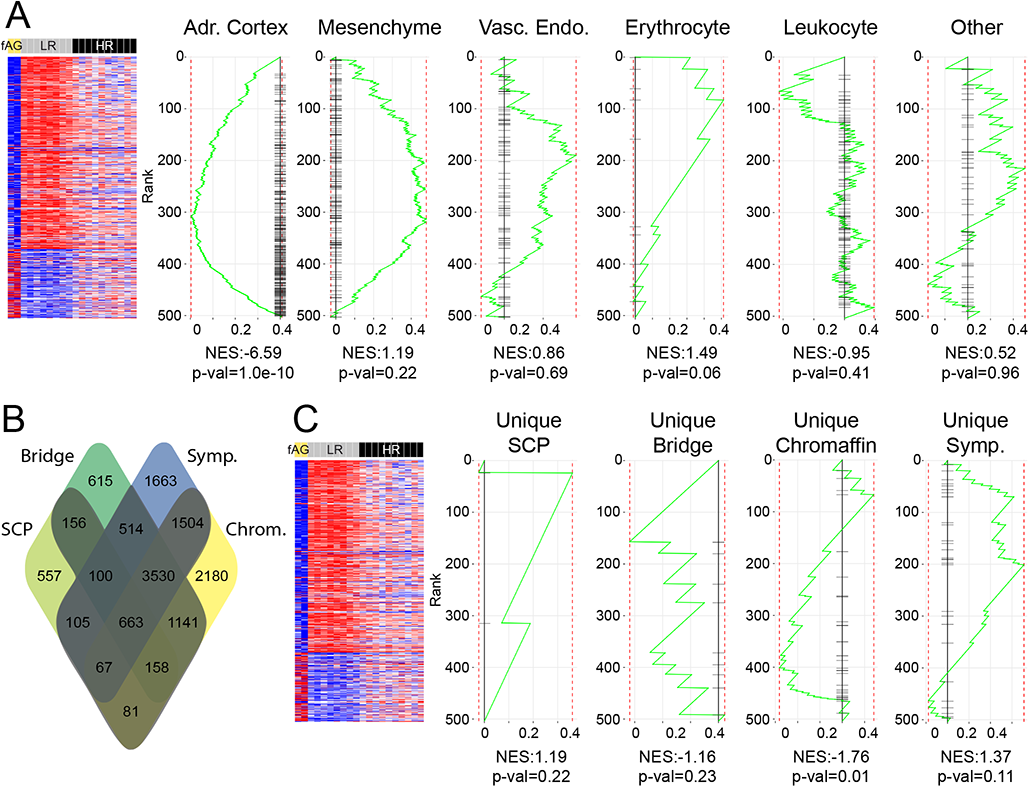
Gene set enrichment analyses of distinct signatures related to human adrenal gland development. (A) Gene set enrichment analysis (GSEA) of the signatures of the non-adrenal medulla cell types identified in the developing human adrenal gland within the list of HR vs LR DEGs, including adrenal cortex, mesenchyme, vascular endothelium, erythrocyte, leukocyte and other. (B) Venn diagram showing the overlapping of the human SCP, highlighting the numbers of genes uniquely associated to SCPs, bridge cells, chromaffin cells or sympathoblasts and (C) their GSEA scores within the list of HR vs LR DEGs.

**Supplemental Figure S3:**
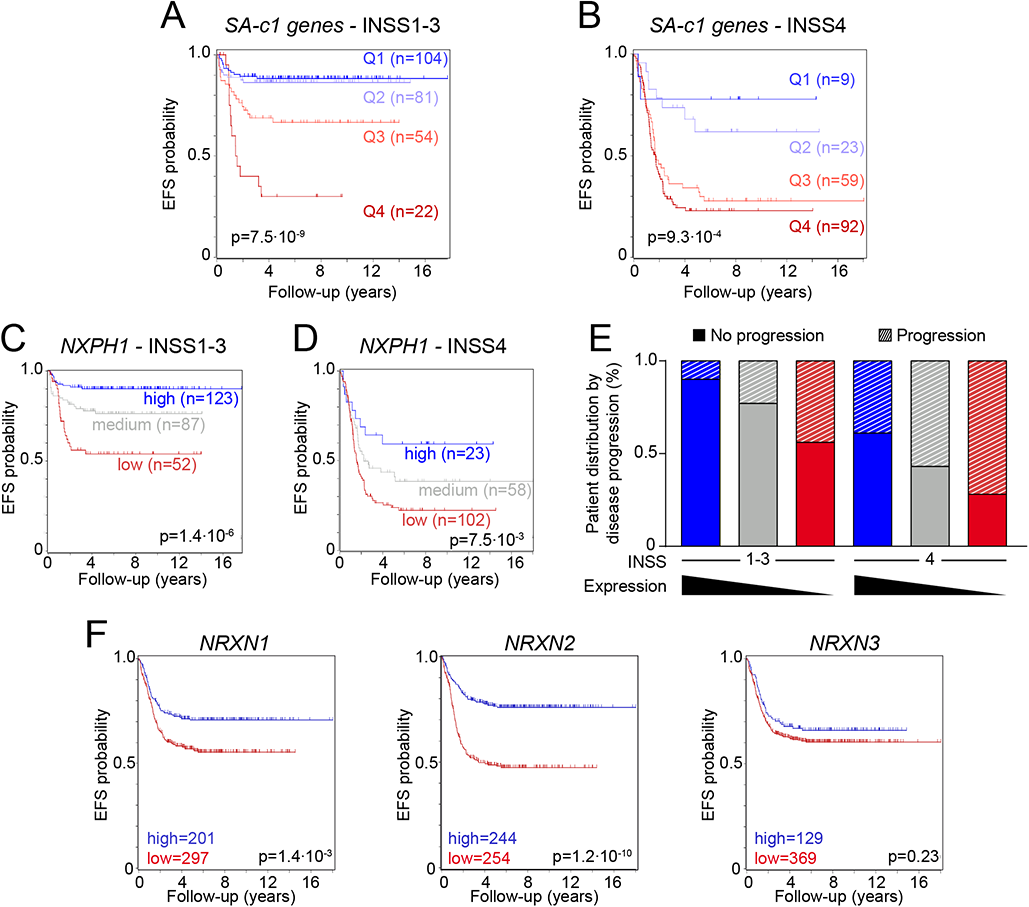
Event-free survival probabilities of NB patient samples based on the expression levels of NXPH1 or NRXN1-3. (A, B) Even-free survival (EFS) probabilities of the SEQC patient samples subdivided into quartiles based on the expression of the SA-c1 genes (Q1-Q4, from higher to lower numbers of SA-c1 genes showing expression levels above average) and INSS tumor stage, subdivided into (A) stages 1-3 and (B) stage 4 sub-groups. (C, D) Even-free survival (EFS) probabilities of the SEQC patient samples based on NXPH1 transcript levels and INSS tumor stage, (C) INSS1-3 and (D) INSS4 sub-groups. (E) Distribution of the sub-groups of INSS stages 1-3 and stage 4 tumor samples based on NXPH1 transcript intensity (high, medium or low) and disease progression. (F) EFS probabilities of the SEQC patient samples based on the expression levels of NRNX1, NRXN2 or NRXN3. Significance was automatically assessed by the R2 server using log-rank test (A-D and F) or with the Chi-square and Fisher’s tests (E).

**Supplemental Figure S4:**
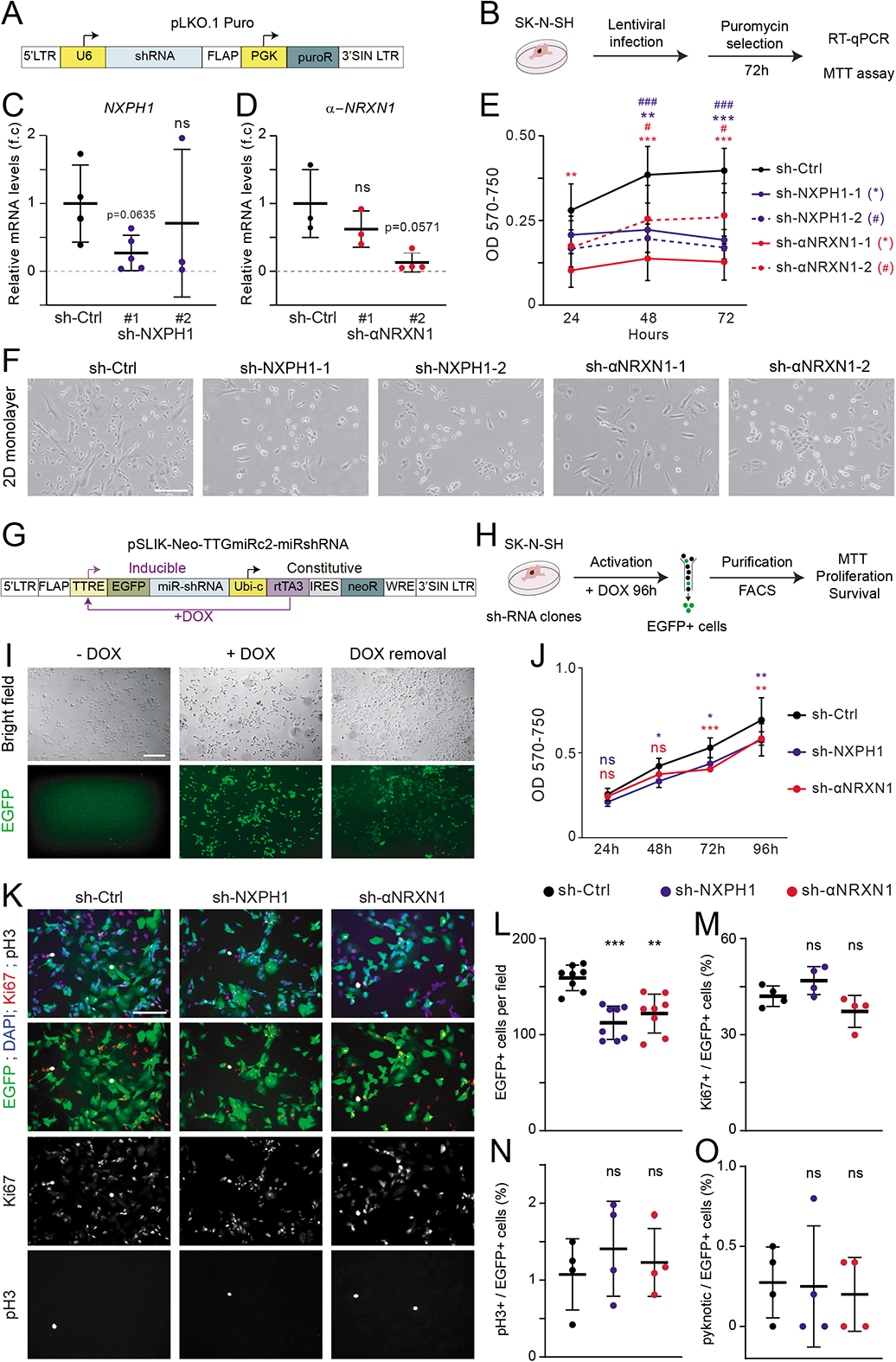
Cellular behaviour of constitutive and inducible sh-NXPH1 and sh-αNRXN1 clones in vitro. (A) Schematic representation of the pLKO.1-Puro construct used to generate stable clones of SK-N-SH cells constitutively expressing sh-RNAs specifically targeting human NXPH1 or α-NRXN1. (B) Experimental design applied in C-F. (C, D) Relative transcripts levels of (C) NXPH1 and (D) α-NRXN1 quantified by RT-qPCR in the sh-Ctrl clone and the two sh-NXPH1 and two sh-αNRXN1 clones generated. The values represent the mean relative mRNA levels ± s.d calculated from 3-4 independent experiments. (E) MTT assay performed over a time-course of 72 hours to assess the cell viability of the sh-Ctrl (black) clone and the two sh-NXPH1 (blue) and two sh-αNRXN1 (red) clones generated. The values represent the mean OD570-750 ± s.d obtained from 4 independent experiments. (F) Bright field images of the sh-Ctrl clone and the two sh-NXPH1 and two sh-αNRXN1 clones, obtained 96 hours after puromycin selection. (G) Schematic representation of the pSLIK-Neo-TTGmiRc2 construct used to generate stable clones of SK-N-SH cells that produce EGFP and sh-RNAs specifically targeting human NXPH1 or α-NRXN1 in an inducible manner upon doxycyline treatment. (H) Experimental design applied in (I-O). (I) Representative images showing how EGFP production is triggered by doxycycline treatment (+DOX, after 2 days) and can be repressed again if doxycycline is removed (-DOX, 3 days later). (J) MTT assay performed over a 96 hours time-course to assess the cell viability of the inducible sh-Ctrl (black), sh-NXPH1 (blue) and sh-αNRXN1 (red) clones. The values represent the mean OD570-750 ± s.d obtained from 2-4 independent experiments. (K) Representative images of the inducible sh-Ctrl (black), sh-NXPH1 (blue) and sh-αNRXN1 (red) clones, their mean number (L) of EGFP+ cells ± s.d quantified per field and their mean proportions of (M) Ki67+;EGFP+/total EGFP+ cells ± s.d, (N) pH3+;EGFP+/total EGFP+ cells ± s.d and (O) pyknotic nuclei EGFP+/total EGFP+ cells ± s.d, quantified 7 days after seeding. Significance was assessed with the non-parametric Mann-Whitney test (C, D) or a 2-way ANOVA + Dunnett’s test (E, J, L-O). *p<0.05, **p<0.01, ***p<0.001, ns: p>0.05. Scale bar: 10µm (K), 100µm (I).

**Supplemental Figure S5:**
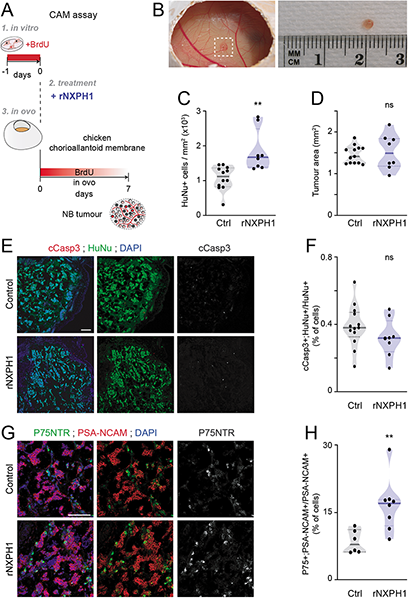
The addition of rNXPH1 stimulates NB growth and increases the proportion of P75/NTR+ NB cells in the CAM assay. (A) Experimental design of the CAM assay used to assess the effects of human recombinant NXPH1 (rNXPH1) on NB cell growth. (B) Representative image of a tumour formed by NB cells 7 days after their seeding onto the CAM. (C) Mean numbers of HuNu+ cells ± s.e.m quantified per area (mm2) in sections of tumours developed by SK-N-SH cells seeded into Matrigel in presence of 10µg/ml rNXPH1 (n=8 tumours) or its control vehicle (BSA, n=13), and (D) the corresponding mean tumour areas ± s.e.m. Each dot represents the value of an individual tumour calculated from 3-6 different confocal images. (E) Representative image showing NB (HuNu+) and apoptotic (cCasp3+) cells and (F) the mean proportion of cCasp3+;HuNu+/total HuNu+ cells ± s.d quantified in sections of tumours developed by SK-N-SH cells seeded into Matrigel in presence of 10µg/ml rNXPH1 (n=8) or its control vehicle (BSA, n=13). (G) Representative image showing NB (PSA-NCAM+) and P75NTR+ cells and (H) the mean proportion of P75NTR+;PSA-NCAM+/total PSA-NCAM+ cells ± s.d quantiified in sections of tumours developed by SK-N-SH cells seeded into Matrigel in presence of 10µg/ml rNXPH1 (n=8) or its control vehicle (BSA, n=6). Each dot represents the value of an individual tumour calculated from 2-6 different confocal images. Significance was assessed with the unpaired t-test (C, F) or the non-parametric Mann-Whitney test (D, H). **p<0.01, ns: p>0.05. Scale bar: 100µm.

**Supplementary Table S1:** DEGs of the LR*vs*fAG and HR*vs*fAG comparisons.

**Supplementary Table S2:** GO terms associated to the LR*vs*fAG and HR*vs*fAG comparisons.

**Supplementary Table S3:** DEGs of the LR*vs*HR comparison.

**Supplementary Table S4:** GO terms associated to clusters c1-c4.

**Supplementary Table S5:** Genes lists corresponding to the distinct sympatho-adrenal signatures.

**Supplementary Table S6:** List of the LR*vs*HR DEGs retrieved in the core SA signature.

